# Wildlife is imperiled in peri-urban landscapes: threats to arboreal mammals

**DOI:** 10.1101/2021.10.30.466631

**Authors:** Oscar M. M Chaves, Júlio Souza Júnior, Gerson Buss, Zelinda MB Hirano, Marcia Jardim, Erica Amaral, João Godoy, Amanda Peruchi, Thais Michel, Júlio César Bicca-Marques

## Abstract

Urbanization and deforestation impose severe challenges to wildlife, particularly for forest-living vertebrates. Understanding how the peri-urban matrix impacts their survival is critical for designing strategies to promote their conservation. We investigated the threats faced by brown howler monkeys (Alouatta guariba clamitans) in peri-urban regions of Rio Grande do Sul (RS) and Santa Catarina (SC) states, southern Brazil, by compiling negative interaction events (hereafter NIE) reported over more than two decades. We assessed the major NIEs, their distribution among age-sex classes, and the predictors of NIE-related mortality. After 20+ years of monitoring, we compiled 540 NIEs (RS = 248 and SC = 292). Electrocution by power lines was the most frequent cause of death or injury (37%), followed by dog attack (34%), vehicle collision (17%), and human mistreatment (12%). The occurrence of lethal injuries ranged from 5% to 69% depending on the type of NIE and on which state it occurred in. The overall post-NIE mortality was 56%. Adults of both sexes were the most affected individuals in both study regions. The minimal adequate GLM model explained 83% of the variation in NIE-related mortality. State, NIE type, and age-sex class were the main predictors of mortality. Overall, mortality was lower in SC and higher among adult females than in the other classes. We found that the survival of brown howler monkeys in the forest-urban interface is constrained by both the urban infrastructure and the growing interactions with humans and domestic and stray dogs (Canis familiaris). We propose the placement of aerial bridges, road signs and speed bumps in areas of frequent animal crossing, the sterilization of stray dogs, and the sensitization of local inhabitants on the importance of respecting and protecting wildlife to reduce their NIEs with humans and domestic animals in the forest-urban interface.

## 1. Introduction

The accelerated destruction of natural habitats by human activities, particularly the expansion of farming, cattle ranching and urbanization (United Nations, 2015; Piano et al., 2020), has resulted in large-scale biodiversity loss (Estrada et al., 2017; Piano et al., 2020). Urban expansion in regions characterized by fragmented landscapes is particularly critical because it imposes additional pressures on threatened species that increase the risk of local extirpation (United Nations, 2015; Salomão et al, 2019; Piano et al., 2020), particularly when the adaptations of the remaining wildlife to the urban landscape increase their encounters with humans (Schell et al., 2020). Therefore, identifying the main threats faced by wildlife in peri-urban landscapes (i.e. mixed landscapes of rural and urban elements that experience intense human pressure: Douglas, 2006) is the first step to developing appropriate conservation strategies aimed at preventing or mitigating their impacts on wild populations.

Urbanization-related processes cause negative impacts on animals worldwide (e.g. butterflies and dung beetles: Salomão et al., 2019; Piano et al., 2020; reptiles: Gonçalves et al., 2018; birds: Bernardino et al., 2018; primates and other mammals: Bueno et al., 2015; Cibot et al., 2015; Katsis et al., 2018; Al-Razi et al., 2019; Galea and Humle, 2021; Jones-Román et al., 2021; wild terrestrial vertebrates in general: Villatoro et al., 2019; Rodríguez et al., 2020; Teixeira et al., 2020). Roads, power lines, houses/buildings, and areas inhabited by domestic and stray dogs increase the risk of death to individuals, particularly to those dispersing through the urban edge or adjacent to it, thereby compromising gene flow between isolated populations immersed in impermeable or semi-permeable urban matrices (Sol et al., 2013; Bernardino et al., 2018; Schell et al., 2020).

Arboreal tropical primates are among the most vulnerable vertebrates to urbanization because of their high dependence on emergent trees (Peres, 1994; Arroyo-Rodríguez and Dias, 2009; Rovero et al., 2015) and because human encroachment into their habitats is increasing (Estrada et al., 2017). While the impact of forest fragmentation, selective logging and hunting on primate behavior and demography has received significant attention (e.g. *Procolobus rufomitratus* and *Colobus guereza*: Gillespie and Chapman, 2008; *Alouatta* spp.: Arroyo-Rodríguez and Dias, 2009; *Ateles geoffroyi*: Chaves et al., 2011; see also Marsh, 2003; Marsh and Chapman, 2013), the impact of urbanization on primate survival in the Neotropics and Afrotropics has been often neglected (but see Gordo et al., 2013; Cibot et al., 2015; Bicca-Marques, 2017; Katsis et al., 2018; Cunneyworth and Duke, 2020; Cunneyworth and Slade, 2021).

Primates inhabiting small habitat patches (i.e. <10 ha; *sensu* Marsh et al., 2003), which may be immersed in peri-urban landscapes, tend to face higher levels of food scarcity, physiological stress and spatial isolation among other adverse consequences of living in these environments (Fahrig, 2003; Arroyo-Rodríguez and Dias, 2009, Bicca-Marques et al., 2020; Cunneyworth and Duke, 2020). Species that cope with these peri-urban stressors can exploit food patches containing native and cultivated plants and human-provisioned or wasted foods in the matrix as have been reported in Africa (e.g. *Papio ursinus*: Beamish and O’Riain, 2004; *Pan troglodytes*: Cibot et al., 2015; *Chlorocebus pygerythrus*: Chapman et al., 2016; *Colobus angolensis* and *Cercopithecus mitis*: Cunneyworth and Slade, 2021) and the Americas (e.g. *Alouatta guariba clamitans*: Chaves and Bicca-Marques, 2017; Corrêa et al., 2018; Back and Bicca-Marques, 2019; *Cebus imitator*: Mckinney, 2011; *Saguinus bicolor*: Gordo et al., 2013). However, peri-urban primates are also exposed to the aforementioned intense vehicle traffic in roads and highways, powerline networks, dog attacks and human mistreatment while navigating between food patches (e.g. Lokschin et al., 2007; Beamish and O’Riain, 2004; Buss, 2012; Gordo et al., 2013; Bicca-Marques, 2017; Bicca-Marques et al., 2020; Azofeifa-Rojas et al., 2021; Cunneyworth and Slade., 2021; Galea and Humle, 2021). Currently, at least 43% of all primate species (or 218 out of 505 spp.) are affected by one or a combination of these urban stressors (Asia = 73 spp., Americas = 65 spp., mainland Africa = 63 spp., Galea and Humle, 2021).

This scenario illustrates the encroached Atlantic Forest landscapes (Ribeiro et al., 2009), where 19 of the 27 nonhuman primates are endemic (Culot et al., 2019), three are Near Threatened, four are Vulnerable, seven are Endangered and five are Critically Endangered (IUCN, 2021). Although the Atlantic Forest is the most developed and populated Brazilian biome (Mittermeier et al., 2004), the impact of urbanization on the conservation status of its threatened primate fauna is poorly known.

The brown howler monkey (*Alouatta guariba clamitans*) is a Vulnerable (Buss et al., 2019) endemic Atlantic Forest primate found in isolated forest patches immersed in peri-urban and rural landscapes of south and southeastern Brazil. The taxon’s ecology and behavior are well-known, particularly in south Brazil (Martins, 2006; Buss, 2012; Chaves and Bicca-Marques, 2013, 2017; Chaves et al., 2018; Corrêa et al., 2018; Back and Bicca-Marques, 2019). However, the lack of long-term data on the influence of peri-urban threats on its populations compromises our assessments of their conservation importance.

In this study we compiled almost three decades of data on NIEs involving free-ranging brown howler monkeys in urban and peri-urban landscapes in the two southernmost Brazilian states (Rio Grande do Sul and Santa Catarina, hereafter RS and SC, respectively). Specifically, we assessed (i) the types of NIE and their relative frequency, (ii) the level of physical harm caused by each NIE, (iii) the proportion of brown howler monkeys that recovered from distinct external injuries and the proportion of those that were released back into their habitats, (iv) the relationship between age-sex class and the frequency of each type of NIE, (v) the role played by season and day of the week on the frequency of NIEs, and (vi) the potential predictors of NIE-related outcomes (i.e. if animals survived or died because of the NIE). Based on the aforementioned, we hypothesized that brown howler monkeys are imperiled in peri-urban areas in both study regions because of the presence of dangerous urban elements such as power lines, roads, and domestic dogs. In light of our findings, we propose management strategies to prevent and reduce the occurrence of NIEs and fatalities involving howler monkeys and other arboreal mammals in peri-urban landscapes.

### 2. Materials and methods

#### 2.1. Study species

Brown howler monkeys, alike their congenerics, are known for their high resilience to habitat disturbance. This resilience has been associated with their highly flexible folivorous-frugivorous diet, including the exploitation of cultivated foods in gardens and orchards (Dias and Rangel-Negrín, 2015; Chaves and Bicca-Marques, 2016, 2017), and their home ranges often <15 ha. Brown howler monkey populations in peri-urban areas in southern Brazil are commonly confined to small (<10 ha) private forest fragments (Printes et al., 2010; Chaves and Bicca-Marques, 2013; Corrêa et al., 2018). These discrete subpopulations may interact as metapopulations and may therefore play an important role in the conservation of this threatened species that is also highly susceptible to outbreaks of yellow fever (Almeida et al., 2012; Bicca-Marques et al., 2017; Buss et al., 2019).

#### 2.2. Study area and forest remnants

In RS, we conducted this study in a ca. 200-km^2^ region in the municipalities of Viamão and Porto Alegre, particularly in urban and peri-urban areas of Viamão and the district of Lami (Fig. 1, Table 1). We focused >90% of our sampling effort in an area of 110 km^2^ (Fig. 1). In SC, we monitored a ca. 800-km^2^ peri-urban region in the municipalities of Blumenau, Indaial, Pomerode and Jaraguá do Sul (Fig. 1, Table 1). Additionally, we occasionally monitored other districts of Porto Alegre, RS, and municipalities along the coastal region of SC when local inhabitants reported NIEs with brown howler monkeys (Fig. 1, Table S1).

**Table 1.** Demographic variables of the main municipalities/cities where NIEs involving brown howler monkeys were monitored in Rio Grande do Sul and Santa Catarina, southern Brazil

| Variable<sup>a</sup> | Rio Grande do Sul |  |  | Santa Catarina<sup>b</sup> |  |  |  |  |
| --- | --- | --- | --- | --- | --- | --- | --- | --- |
|  | Viamão | Lami | RS | BL | IN | PO | JS | SC |
| Area (km<sup>2</sup>) | 1,496 | 28.2 | 281,707 | 518.6 | 430.8 | 214.3 | 530.1 | 95,731 |
| Population size in 2019 | 252,872 | 4,642 | 11,377,239 | 357,199 | 69,425 | 33,447 | 177,697 | 7,164,788 |
| Urban population | 224,943 | — | 9,100,291 | 294,773 | 52,927 | 23,823 | 132,800 | 5,247,913 |
| Rural population | 14,441 | — | 1,593,64 | 14,238 | 1,927 | 3,936 | 10,323 | 1,000,523 |
| % population in urban areas | 94.0 | — | 99.8 | 95.4 | 96.5 | 85.8 | 92.8 | 84.0 |
| Population density (ind./km<sup>2</sup>) | 160 | 165 | 39.8 | 596.1 | 127.3 | 129.3 | 270.3 | 65.3 |
| Population growth (%) | 5.3 | — | 20.0 | 13.5 | 21.0 | 17.0 | 19.5 | 12.8 |
| #vehicles in 2018 | 95,734 | — | 5,365,382 | 197,586 | 33,894 | 18,100 | 84,776 | 3,672,593 |
| #Urban residences | 70,514 | 4,030 | 3,084,215 | 96,866 | 16,753 | 7,423 | 42,070 | 1,691,822 |
| #Rural residences | 4,883 | — | 515,589 | 4,196 | 614 | 1,130 | 3,036 | 301,190 |

<sup>a</sup>Information sources were: Instituto Brasileiro de Geografia e Estatística (population census 2010; IBGE, 2020) and Departamento Nacional de Trânsito (number of vehicles 2018; Denatran, 2020). Population growth was based on the last 10 years (i.e. the differences between the population estimate of 2019 and the population census of 2010). Vehicles included: cars, pickups, trucks, buses, and minibuses.
<sup>b</sup>Municipality abbreviations: BL=Blumenau, IN= Indaial, PO=Pomerode, and JS=Jaraguá do Sul.
— Information not available.

**Fig. 1.**
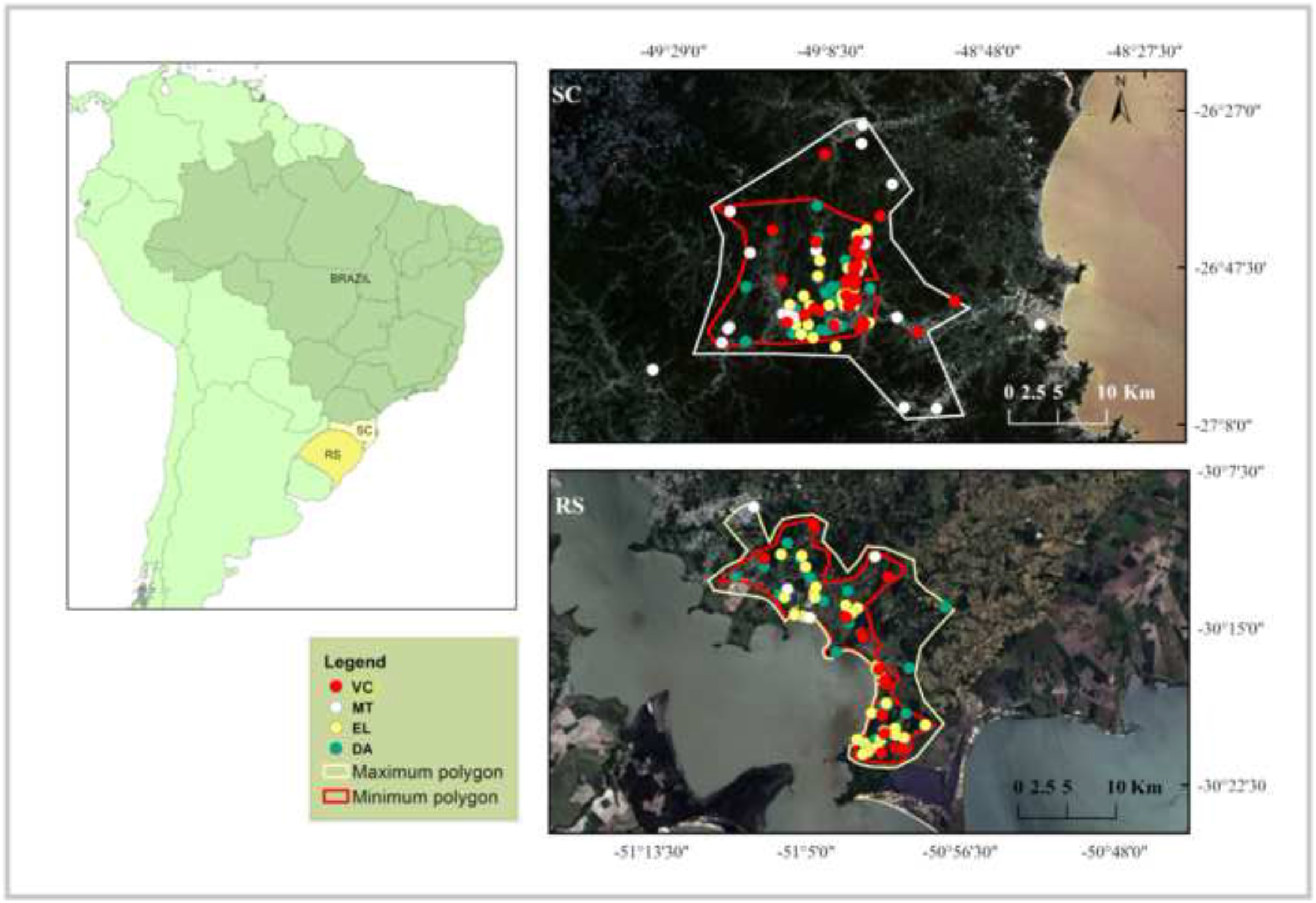
Location of NIEs involving brown howler monkeys in Santa Catarina and Rio Grande do Sul states, southern Brazil. Color circles represent each type of NIE. The white polygon includes >98% of NIEs, while the red polygon includes the study region with higher density of NIEs (excluding outliers). Free open-access images (available at http://www.cbers.inpe.br//) from 22 November 2021.

Human populations grew as little as 5% in Viamão to as much as 21% in Indaial from 2000 to 2010, reaching densities ranging from 160 people/km^2^ in Viamão to 596 people/km^2^ in Blumenau (Table 1). Most people (≥86%) live in peri-urban areas in the study regions. The number of houses vary from about 4,000 in Lami to 97,000 in Blumenau (IBGE, 2020; Table 1). Most of the study areas are surrounded by <1-ha to 100-ha Atlantic Forest fragments and scattered vegetation corridors in different successional stages. Subtropical semideciduous forests dominate the vegetation in both study regions.

Regardless of fragment size and level of official environmental protection, brown howler monkeys that move between habitat patches in these peri-urban landscapes face risks imposed by roads, power lines, and human settlements (Table 1). These structural elements together with pastures and cultivated lands reduce matrix permeability, compromising inter-path connectivity. Despite these threats for moving and dispersing individuals and the human pressures on the plant community structure of habitat patches (e.g. selective logging, residential development), brown howler monkey populations have persisted.

#### 2.3. Data collection

We recorded the NIEs involving brown howler monkeys between 1995 and 2021 in RS and between 1991 and 2020 in SC (Fig. S1). We used four sources of information: (i) our own field observations, and reports from local (ii) inhabitants, (iii) environmental authorities (i.e. municipal and state environmental secretariats, Environmental Military Police /SC), and (iv) wildlife rehabilitation centers and veterinary hospitals and clinics (Porto Alegre and Viamão, RS). We visited the location of ∼70% of reported NIEs to record the following information: geographic coordinates using a Garmin GPS, type of NIE (electrocution or sub-lethal injuries in power lines, EL; vehicle collision with any kind of motor-vehicle, VC; domestic dog attack, DA; and human mistreatment (including illegal captivity and physical mistreatments, MT; Fig. 2), external injury level (mild-medium, severe, or lethal) (Fig. S2), and, whenever possible, the fate of the injured individual. Type MT involved firearm shooting, stoning, and illegal captivity. The last is associated with chaining, inadequate feeding, precarious sanitary conditions, and lack of veterinarian care.

**Fig. 2.** Main threats faced by brown howler monkeys (*Alouatta guariba clamitans*) in urban and peri-urban areas in Rio Grande do Sul state, southern Brazil. Adult female using a power line to cross a road (A), individuals electrocuted and/or mutilated on power lines (B-D), monkeys crossing roads to access food patches (E, F), adult and sub-adult males vehicle collision (G, H), adult male on cultivated tree in a subsistence orchard guarded by dogs (I), juvenile individual walking on the ground near a domestic dog (J), victim of mistreatment in a peri-urban area of RS (K). Photos by Ó. M. Chaves (A-C, H), J. C. Godoy (D-F), G. Buss (G, K) and J. P. Back (I-J).

The mild-medium injury level of brown howler monkeys included minor scratches that did not require prompt veterinarian care (e.g. slight skin-burns and teeth loss) and injuries that required surgery (e.g. bleeding, multiple dog bites, bone fractures, amputation of fingers, limbs or tail; Fig. S2). Severe injuries included multiple wounds that could lead to death without urgent veterinarian intervention (Fig. S2). These injuries often impeded the release of the individual back into the wild upon its recovery. Finally, lethal injuries often caused the howler’s death up to 5 h after the NIE.

Whenever possible, we frozen the brown howler monkey carcasses in the collection of biological material of CEPESBI in SC, and in the Laboratório de Primatologia or the Museu de Ciências e Tecnologia/PUCRS, or the Museu de Ciências Naturais (SEMA/RS) in RS. A subsample of carcasses from RS was necropsied in a study on the taxon’s helminth parasite fauna (Lopes et al., 2021). Injured brown howler monkeys were rescued by local authorities, researchers, or volunteers, who, then, sent them to veterinarian hospitals/clinics or authorized wildlife rehabilitation centers. The full NIE dataset is available in Chaves et al. (2021).

#### 2.4. Database limitations

Although we recorded NIEs involving brown howler monkeys during almost three decades in each study region, we are conservative in extrapolating and interpreting our findings because of several imitations inherent of this kind of long-term study. We identified five major limitations that may have influenced the patterns that we found. First, we certainly missed NIEs (Fig. 1) that were not detected, reported by local people or not forwarded to us by local authorities. This situation is more likely when the injuries were mild-medium and when the monkey returned to its group soon after the NIE (Óscar M. Chaves, personal observation). Second, our sampling effort varied over time (Fig. S1) given temporal changes in the number of researchers, volunteers, and local informants. In this respect, the 1990s were poorly sampled because of a lack of volunteers or institutional groups to rescue the brown howler monkeys.

Third, the interest of local people in reporting NIEs may vary over time and between study regions, compromising the standardization of sampling effort. While there is a long-term, well-consolidated project (Projeto Bugio-FURB) monitoring NIEs in SC that provides veterinarian care to injured animals, and that promotes the participation of local inhabitants, a similar interinstitutional effort is incipient in RS. Fourth, given the large sampling areas in both study regions (see Fig. 1) and the lack of reliable data on the size of their brown howler monkey populations, we could not estimate the proportion of individuals dying after NIEs. Finally, local environmental authorities were more active collaborators in SC than in RS. This difference may explain the greater number of records of human mistreatment in SC. Despite these and other limitations, our database represents a useful description of the main threats faced by brown howler monkeys living in the forest-urban interface for promoting their conservation via the design of appropriate management strategies.

#### 2.5. Characterization of the study regions

We estimated 10 structural variables of the peri-urban matrices for those NIEs for which we have precise geographic coordinates, date of occurrence, type of NIE, and injury level (*n* = 335, 212 in SC and 123 in RS, see Table 2) to assess their relationship with NIE lethality (i.e. the probability of an individual to die from a particular NIE): (1) matrix element where the NIE occurred, (2) NIE type, (3) number of houses within a 500-m radius from the location of the NIE, (4) total number of elements in the peri-urban matrix (e.g. roads, houses, buildings, airports, power lines, gardens, orchards, pastures, and others) within a 500-m radius from the location of the NIE, (5) type of road (primary or secondary), (6) road material (paved or unpaved), (7) distance to the nearest road, (8) distance to the nearest small forest fragment <10 ha, (9) distance to the nearest ≥70 ha-forest fragment, and (10) distance to the nearest house. We estimated these traits by exporting the Global Positioning System (GPS) locations of the NIEs from the software Map Source 6.16.3 (Garmin^®^) to Google Earth Pro (Google^®^). We chose a high-resolution satellite image (with a low percentage of clouds and shadows) of the year of the NIE for each GPS position using the option ‘historic images,’ which includes images from 2002 to 2019. We analyzed Landsat 5 images in the software QGIS 3.6 to estimate the variables for those NIEs that occurred between 1991 and 2001 (*n =* 46).

**Table 2.** Potential predictors of post-NIE lethality in brown howler monkeys in the peri-urban matrices of Rio Grande do Sul and Santa Catarina states, southern Brazil

| Predictor<sup>a</sup> | Description | Effect<sup>b</sup> |
| --- | --- | --- |
| *Structure of peri-urban matrix* |  |  |
| **1) Matrix element** | Element of the urban matrix where the NIE occurred, including roads, gardens, cities, fragment edges, forest remnants, etc. | (+) |
| **2) NIE type** | Main NIE involving brown howler monkeys: electrocution (EL), dog attack (DA), vehicle collision (VC), and mistreatment (MT) | N.A. |
| **3) # houses** | Number of houses or clearly identifiable roofs around a 500-m radius from the NIE location | (+) |
| **4) # elements** | Total number of elements constituting the anthropogenic matrix (e.g. roads, airports, power lines, houses, buildings, gardens, and pastures) around a radius of 500 m from the NIE location | (+) |
| **5) Type of road** | Type of road nearest to the NIE location: primary large road (>15 m wide) with high vehicle traffic (P), and secondary small roads (<10 m wide) with low vehicle traffic (S) | P>S |
| **6) Road material** | If the road was paved or unpaved | paved>unpaved |
| **7) DNR** | Distance from the NIE location to the nearest primary or secondary road (m) | (-) |
| **8) DNS** | Distance from the NIE location to the nearest small fragment <10 ha | (+) |
| **9) DNL** | Distance from the NIE location to the nearest large fragment > 80 ha | (+) |
| **10) DNH** | Distance from the NIE location to the nearest house | (-) |
| *Other factors* |  |  |
| 11) Age-sex | Age-sex class of the individual, including adults (A), subadults (S), and juveniles (J) of both sexes. | N.A. |
| 12) Study region | The study was performed in two Brazilian states: Santa Catarina (SC) and Rio Grande do Sul (RS) | N.A. |
| 13) Season | Season of the year in which each NIE occurred: summer (Su), fall (F), winter (W), and spring (Sp) | >S |
| 14) Day | Day of the week in which each NIE occurred: Monday (Mon), Tuesday (Tue), Wednesday (Wed), Thursday (Thu), Friday (Fri), Saturday (Sat), and Sunday (Sun) | >Sat/Sun |
| 15) Incident*element | Interaction between the type of NIE and the matrix element | N.A. |
| 16) Incident*age-sex | Interaction between the type of NIE and the age-sex category | N.A. |

<sup>a</sup>Ten physical characteristics of the peri-urban matrices estimated in this study enhanced in bold.
<sup>b</sup>Expected effect according to available evidence: positive (+), negative (-), higher (>), and non-assessed (N.A.).

#### 2.6. Statistical analyses

We performed Chi-square tests for proportions using the R function ‘prop.test’ to compare the proportion of occurrence of each type of NIE involving brown howler monkeys, age-sex classes, seasons, and days of the week. When we found significant differences, we compared the proportion of records in each variable via post-hoc proportion contrasts using the R function ‘pairwise.prop.test’ with a Bonferroni correction. We performed generalized linear mixed models (GLMM; Zuur et al., 2009) using the function ‘lmer’ of the R package lme4 to assess the influence of the 16 predictor variables listed in Table 2 on NIE-related mortality. We set the binomial family error for the response variable (i.e. if individuals died or survived following the NIE) and a log link for running the models. We specified the 16 variables as fixed factors and the sampled year-ID as random factor to account for repeated-measures during the same years. We considered only two second-order interactions that are ecologically relevant, namely NIE type*matrix element and NIE type*age-sex class, to minimize overparameterization and problems of convergence of the global model (the model containing all fixed and random factors) due to the inclusion of a large number of variables and their interactions (Grueber et al., 2011). Before running this analysis we tested the variables for multicollinearity using the ‘vifstep’ function of R package dplyr. We included all variables in the global model because their Variance Inflation Factors (VIFs) were <3.

Then, we used the model simplification procedure to determine the minimal adequate (most ‘parsimonious’) model. In this method, the maximal model is simplified over a backward stepwise procedure until a model that produces the least unexplained variation or the lowest Akaike’s Information Criterion (AIC) is found (Crawley, 2012). We used the AICc to select the ‘best’ model as recommended when sample size/number of predictor variables <40 (Burnham and Anderson, 2003). We used a likelihood ratio test over the R function ‘anova’ to test the significance of the ‘best model’ in comparison with the null model (the model including only the random factor). Finally, we used the ‘r.squaredGLMM’ function of the R package MuMIn (Barton, 2016) to estimate an equivalent of the coefficient of determination or pseudo-R^2^ for the ‘best’ GLMM. The datasets used to perform these analyses are available in Chaves et al. (2021). We ran all statistical analyses in R v.3.6.3 (R CoreTeam, 2020).

## 3. Results

### 3.1. Major NIEs involving brown howler monkeys in peri-urban matrices

We recorded 540 NIEs involving brown howler monkeys in the peri-urban matrices of RS (*n* = 248) and SC (*n* = 292), from which we discarded 56 from further analysis because of incomplete information on the date, NIE type and/or injury level. Then, we collected complete information for 484 NIEs. In addition to our main study regions, we included NIEs in other 11 municipalities in RS and 24 in SC (6% and 22% of state’s NIEs, respectively, Table S1). The major NIEs were electrocution (37% of 488 NIEs with complete information), followed by dog attack (34%), vehicle collision (17%), and human mistreatment (12%, Figs. 2 and 3 A-C). The vast majority of NIEs occurred at daytime when brown howler monkeys walked on power lines (Fig. 2 A-D), tried to cross paved or unpaved roads (Fig. 2 E-H), or descended to the ground (Fig. 2 I-J) to cross canopy gaps or to move between forest patches. A high percentage of the dog attacks (66% in RS and 49% in SC; Fig. 3 A, C) were lethal. On no occasion did the killer dogs eat the monkey’s flesh. Dog attacks involved stray and domestic dogs, and in all cases, they abandoned the carcass *in situ* upon the monkey’s death. Furthermore, brown howler monkeys kept illegally in captivity, commonly infants and juveniles, represented most records of human mistreatment (87% of 60 NIEs). The remaining cases were brown howler monkeys shot with ball-bearing guns by local inhabitants.

**Fig. 3.**
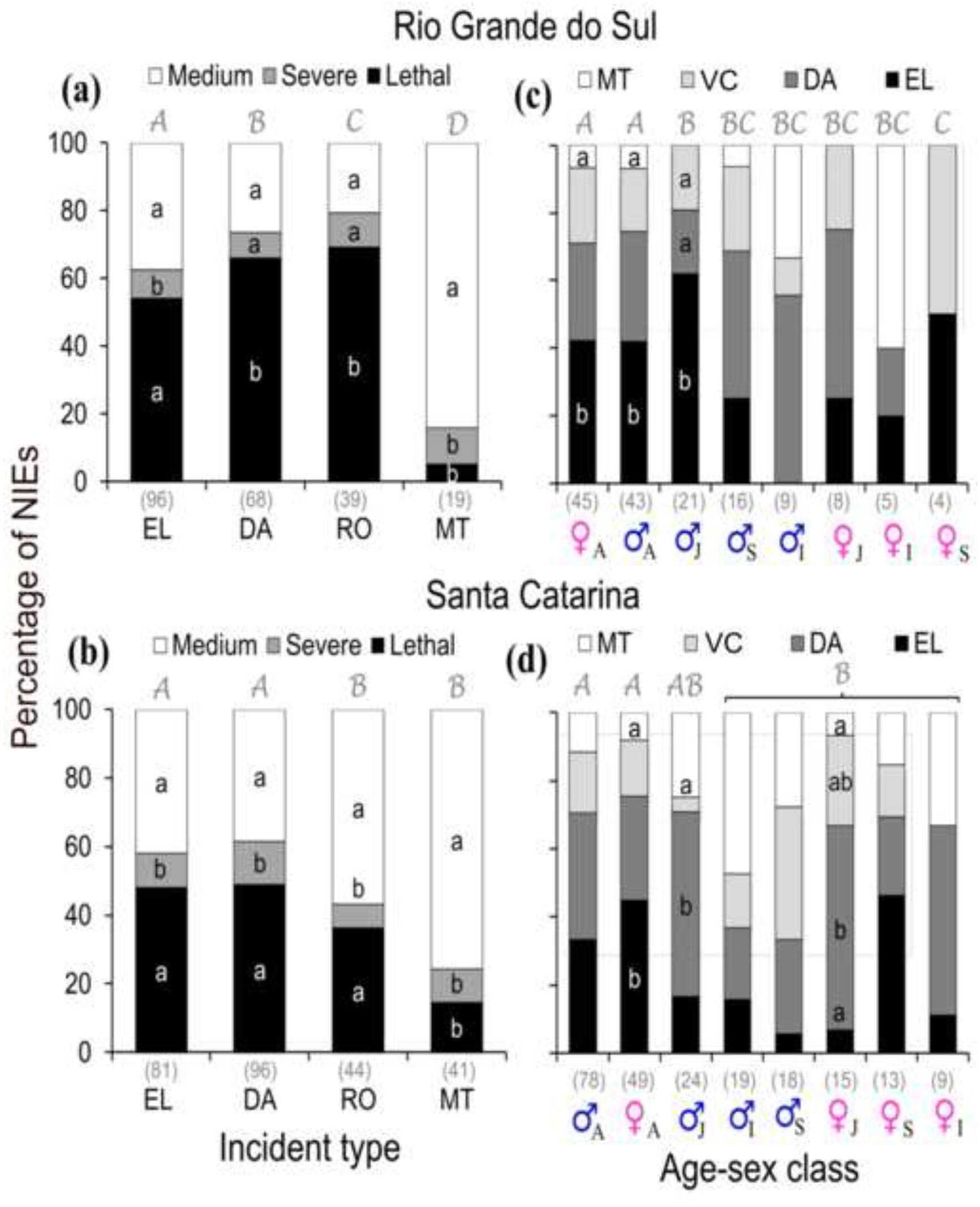
Comparison of the proportion of NIEs involving brown howler monkeys according to the type of incident (a, b) and the age-sex class (c, d) in the States of Rio Grande do Sul (top panels) and Santa Catarina (bottom panels). Different Lucida handwriting capital letters on the bars indicate differences among incident types or age-sex classes, and lowercase letters inside the bars indicate differences among injury levels or incident types (proportion contrasts, *P* < 0.05). When no significant differences were detected within each incident type or age-sex class (proportion contrasts, *P* > 0.05), no lowercase is show. Type of incident: EL = electrocution, DA = dog attack, RO= vehicle collision, and MT= human mistreatment (further details in Methods). Numbers in parentheses at the bottom of bars represent the number of events per type of NIE or age-sex class. Age categories: A = adult, S= subadult, J = juvenile, and I = infant. Total number of NIEs considered in each graph: 222 (a), 151 (b), 262 (c), and 225 (d).

The frequency of each type of NIE involving brown howler monkeys varied between RS and SC. The number of records also differed among NIE types in RS (EL = 43% of 222 NIEs, DA = 31%, VC = 18%, and MT = 8%; *χ*^*2*^ = 81, d.f. = 3, *P* < 0.0001; contrasts, *P* < 0.05 in all significant comparisons; Fig. 3 A) and SC (DA = 37% of 262 NIEs, EL = 31%, VC = 17%, and MT = 15%; *χ*^*2*^ = 45, d.f. = 3, *P* < 0.0001, contrasts, *P* < 0.05 in all significant comparisons, Fig. 3 B).

### 3.2. Injury level in brown howler monkeys during NIEs

Most RS brown howler monkeys involved in EL (54%), DA (66%) and VC (69%) suffered lethal injuries (contrasts, *P* < 0.05 in all significant comparisons). The remaining individuals survived with mild-medium (38%, 26%, and 21%, respectively) or severe injuries (8%, 7%, and 10%, respectively; Fig. 3 A). A higher proportion of the individuals involved in EL suffered lethal or mild-medium injuries than severe injuries, while a higher proportion of those involved in DA and VC suffered lethal than mild-medium or severe injuries (Fig. 3 A, contrasts, *P* < 0.05 in all significant comparisons).

Lethal injuries were less frequent in SC brown howler monkeys. They ranged from ca. 10% in MT to 49% in DA (Fig. 3 B). The other individuals involved in these NIEs survived with mild-medium (42%, 39%, and 57%, respectively) or severe injuries (10%, 13%, and 7%, respectively; Fig. 3 B). The proportion of victims of these NIEs with lethal or mild-medium injuries was higher than the proportion with severe injuries (Fig. 3 B, contrasts, *P* < 0.05 in all significant comparisons). A higher proportion of brown howler monkeys involved in MT suffered mild-medium (RS = 11%. SC = 10%) than severe or lethal injuries (RS = 5%, SC = 15%; Fig. 3 A, B, contrasts, *P* < 0.05 in all significant comparisons).

Finally, 56% (269 out of 484 NIEs, Table S2) of the brown howler monkeys with lethal injuries or with mild-medium or severe injuries that were alive immediately following the NIE died after <1 to 8 h during the transport to the veterinarian clinic or during the emergency veterinarian care. The health problems associated with their deaths included cardiorespiratory problems, lung perforations, internal hemorrhages, myases, and mutilations. This mortality represented 61% and 51% of the total number of NIEs with complete data reported for RS and SC, respectively (Table S2). Injured and/or mutilated survivors that were kept for life in public or private wildlife rescue centers represented 25% (RS) and 15% (SC), whereas individuals released back into their habitats summed only 7% (RS) and 2% (SC). The fate of the remaining survivors is unknown.

### 3.3. NIE distribution among age-sex classes

NIEs involving brown howler monkeys affected all age-sex classes in both study regions with a bias towards adult males and adult females (RS: *χ*^*2*^ = 115, d.f. = 7, *P* < 0.0001; SC: *χ*^*2*^ = 158, d.f. = 7, *P <* 0.0001; contrasts, *P*<0.05 in all significant comparisons; Fig. 3 C, D). The proportion of records per NIE type was often similar within each age-sex class (Fig. 3 C, D). The exceptions were higher proportions of EL than MT records for adults of both sexes in RS and for adult females in SC. In SC, juvenile males were more impacted by DA than by VC, and juvenile females were more impacted by DA than by EL and MT (contrasts, *P* < 0.05 in all significant comparisons, Fig. 3 D).

### 3.4. Temporal patterns in the number of NIEs involving brown howler monkeys

The average number of NIEs involving brown howler monkeys per year (mean ± SD) was similar between RS and SC (13 ± 8 *vs* 12 ± 9 NIEs, respectively, Fig. S1). There was a higher frequency of NIEs in the summer and fall than in the winter in RS (*χ*^*2*^ = 30, d.f. = 3, *P* < 0.0001; contrasts, *P*<0.05 in all significant comparisons, Fig. 4 A). The number of NIEs also differed among months in each season (*χ*^*2*^ ranged from 16 to 57, d.f. ranged from 3 to 4 in all cases, *P* < 0.001 in all cases; Fig. 4 A). The month with the greatest number of NIEs in summer, fall, winter, and spring were, respectively, March, April and May, September, and October (contrasts, *P* < 0.05 in all significant comparisons, Fig. 4 A). In contrast, the frequency of NIEs in SC was similar in all seasons (*χ*^*2*^ = 5, d.f. = 3, *P* = 0.2, Fig. 4 B). However, the number of NIEs also differed among months in each season (*χ*^*2*^ ranged from 15 to 23, d.f. ranged from 3 to 4, *P* < 0.005 in all cases; Fig. 4 B), and the month with the highest number of NIEs in summer, fall, winter, and spring were, respectively, January, April, August, and November (contrasts, *P* < 0.05 in all significant comparisons, Fig. 4 B).

**Fig. 4.**
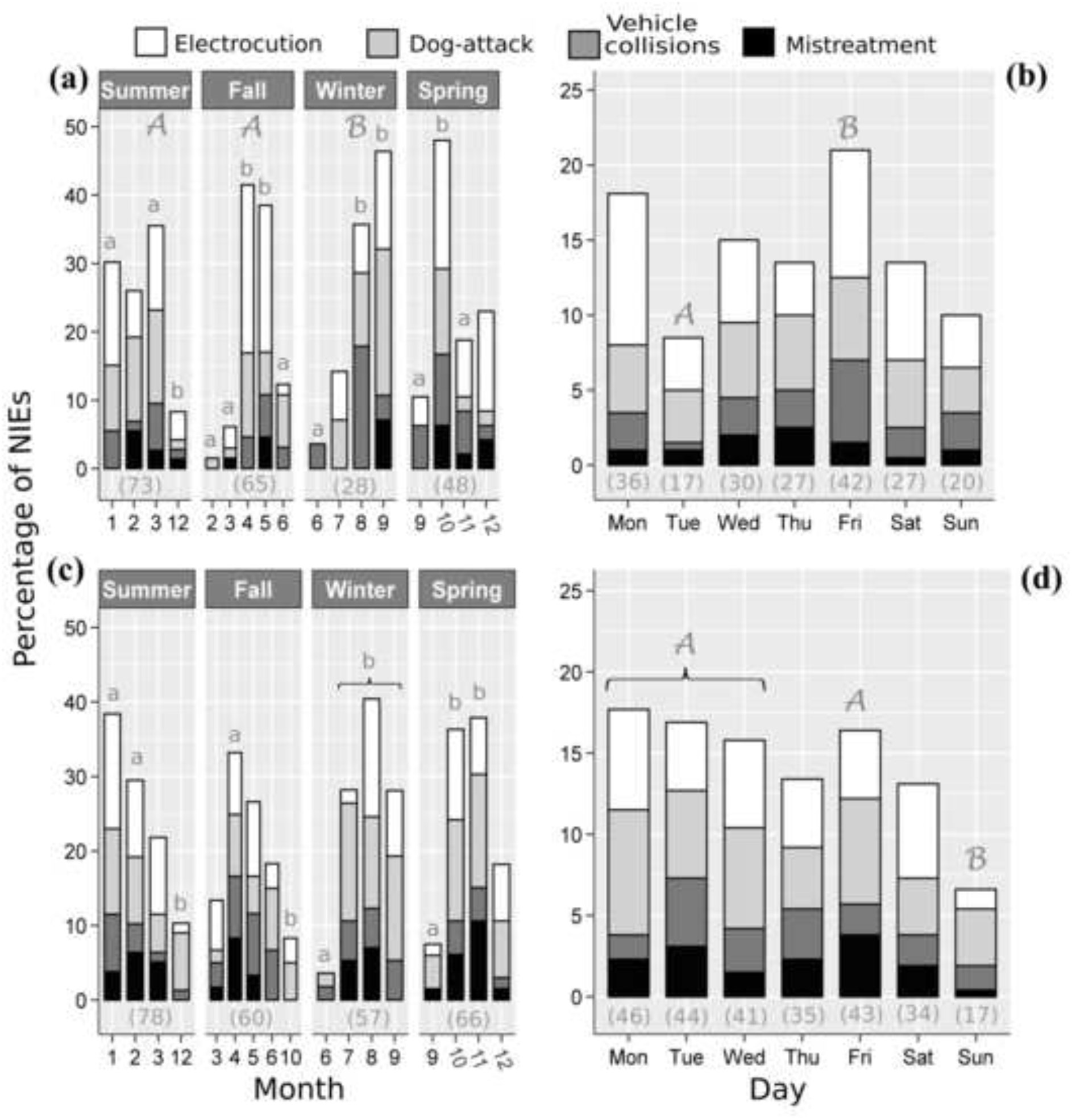
Temporal patterns in the proportion of NIEs according to season (a, b) and day of the week (c, d) in Rio Grande do Sul (top panels) and Santa Catarina (bottom panels) states. Different Lucida handwriting capital letters on the bars indicate differences among seasons or day of the week (proportion contrasts, *P* < 0.05). When the proportion of NIEs was similar (proportion contrasts, *P* > 0.05) among seasons or days, no capital letter is shown. Lowercase letters on the bars in (a) and (b) indicate differences among months within each season (contrasts, *P* < 0.05). When the proportion of NIEs was similar between months (contrasts, *P* > 0.05), no lowercase is shown. Days: Monday (Mon), Tuesday (Tue), Wednesday (Wed), Thursday (Thu), Friday (Fri), Saturday (Sat), and Sunday (Sun). Numbers in parentheses at the bottom of bars represent the number of NIEs per season or day recorded until April 2021. Total number of NIEs considered in each graph: 214 (a), 199 (b), 261 (c), and 260 (d).

The frequency of NIEs involving brown howler monkeys also differed between the days of the week in RS (*χ*^*2*^ = 18, d.f. = 6, *P* = 0.005), because of a greater number of reports of NIEs on Fridays than on Tuesdays (proportion contrast, *P* < 0.05; Fig. 4 C). We also found differences in the frequency of NIEs in SC (*χ*^*2*^ = 19, d.f. = 6, *P* = 0.005, Fig. 4 D). with a greater number of NIEs occurring on Mondays, Tuesdays, Wednesdays and Fridays than on Sundays (contrasts, *P* < 0.05 all significant comparisons; Fig. 4 D).

### 3.5. Predictors of NIE lethality

The minimal adequate GLMM explained 83% of the variation in the lethality of NIEs involving brown howler monkeys and included the predictors ‘study region’, ‘type of NIE’, ‘age-sex class’, ‘day of the week’, and ‘distance to the nearest large forest fragment’ (*R*^*2*^_*c*_ = 0.83; Table 3). NIE-related mortality was lower in SC than in RS (β = -1.2, z-value = -3, *P* < 0.01) and for MT than for the other NIEs (β = -1.8, z-value = -3, *P* < 0.01). Lethality was higher for adult females than for individuals belonging to other age-sex classes (β = 1.9, z-value = 2, *P*<0.05) and for Tuesday NIEs than for those occurring in the other days (β = -1.2, z-value = 2, *P* < 0.05; Table 3).

**Table 3.** Minimal adequate GLMM predicting the lethality of NIE involving brown howler monkeys in the peri-urban anthropogenic matrices of southern Brazil

| Predictors | Parameters<sup>a</sup> |  |  |  |  |
| --- | --- | --- | --- | --- | --- |
| | $\beta$ | SE | z-value | AICc | $R^2_c$ |
| **Model: study region+NIE+age-sex+day+dlf** |  |  | 6.2** | 320 | 0.83 |
| *Study region* |  |  |  |  |  |
| Santa Catarina | -1.19 | 0.36 | -3.3** |  |  |
| *Type of NIE* |  |  |  |  |  |
| Mistreatment | -1.76 | 0.57 | -3.1** |  |  |
| Vehicle collision | -0.61 | 0.44 | -1.4 |  |  |
| Electrocution | -0.30 | 0.39 | -0.8 |  |  |
| *Age-sex class (age-sex)* |  |  |  |  |  |
| Adult female | 1.92 | 0.93 | 2.1* |  |  |
| Subadult male | 1.66 | 1.10 | 1.6 |  |  |
| Adult male | 1.32 | 0.87 | 1.5 |  |  |
| Juvenile female | 1.44 | 1.06 | 1.4 |  |  |
| *Day of the week (day)* |  |  |  |  |  |
| Tuesday | 1.19 | 0.59 | 2.1* |  |  |
| Thursday | 0.72 | 0.57 | 1.3 |  |  |
| *Distance to the nearest large forest (dlf)* | -0.00 | 0.00 | -0.9 |  |  |

<sup>a</sup>Parameters shown: partial regression coefficients ( $\beta_i$ ), standard errors that incorporate model uncertainty (SE), Akaike's Information Criterion for small samples (AICc), and pseudo- $R^2$ ( $R^2_c$ ) indicating the percentage of the variance explained by the fixed and random factors in the minimal adequate model. Significance level: \* $P < 0.05$ , \*\* $P < 0.01$ .

## 4. Discussion

In this study we present an important scientific diagnostic of the major threats faced by brown howler monkeys in peri-urban landscapes of southern Brazil. We found that electrocution was the most frequent NIE affecting the physical integrity of brown howler monkeys alike reported for wildlife worldwide. Power lines kill hundreds of primates (e.g. *Alouatta guariba* c*lamitans*: Lokschin et al., 2007; *Colobus angolensis, Cercopithecus mitis*, and *Otolemur garnettii*: Katsis et al., 2018; *Macaca sinica*: Dittus, 2020; 8 spp. in the Americas, 16 spp. in Africa and 23 spp. in Asia: Galea and Humle, 2021; *Alouatta palliata*: Azofeifa-Rojas et al., 2021; Jones-Román et al., 2021) and hundreds of thousands to millions of birds and other vertebrates each year (Bernardino et al., 2018; Biasotto and Kindel, 2018). Therefore, the implementation of management strategies including the trimming of tree branches, insulation of powerlines, installation of wildlife crossings (i.e. canopy-to-canopy aerial bridges), and an efficient protection of biological corridors (e.g. live fences with native trees) are urgent not only to prevent the electrocution of arboreal wildlife, but to increase habitat connectivity and gene flow between animal populations (Table 4). Similar strategies have been suggested to avoid the electrocution of primates in peri-urban African and Asian landscapes (e.g. Katsis et al., 2018; Al-Razi et al., 2019; Cunneyworth and Slade, 2021; Galea and Humle, 2021).

**Table 4.**
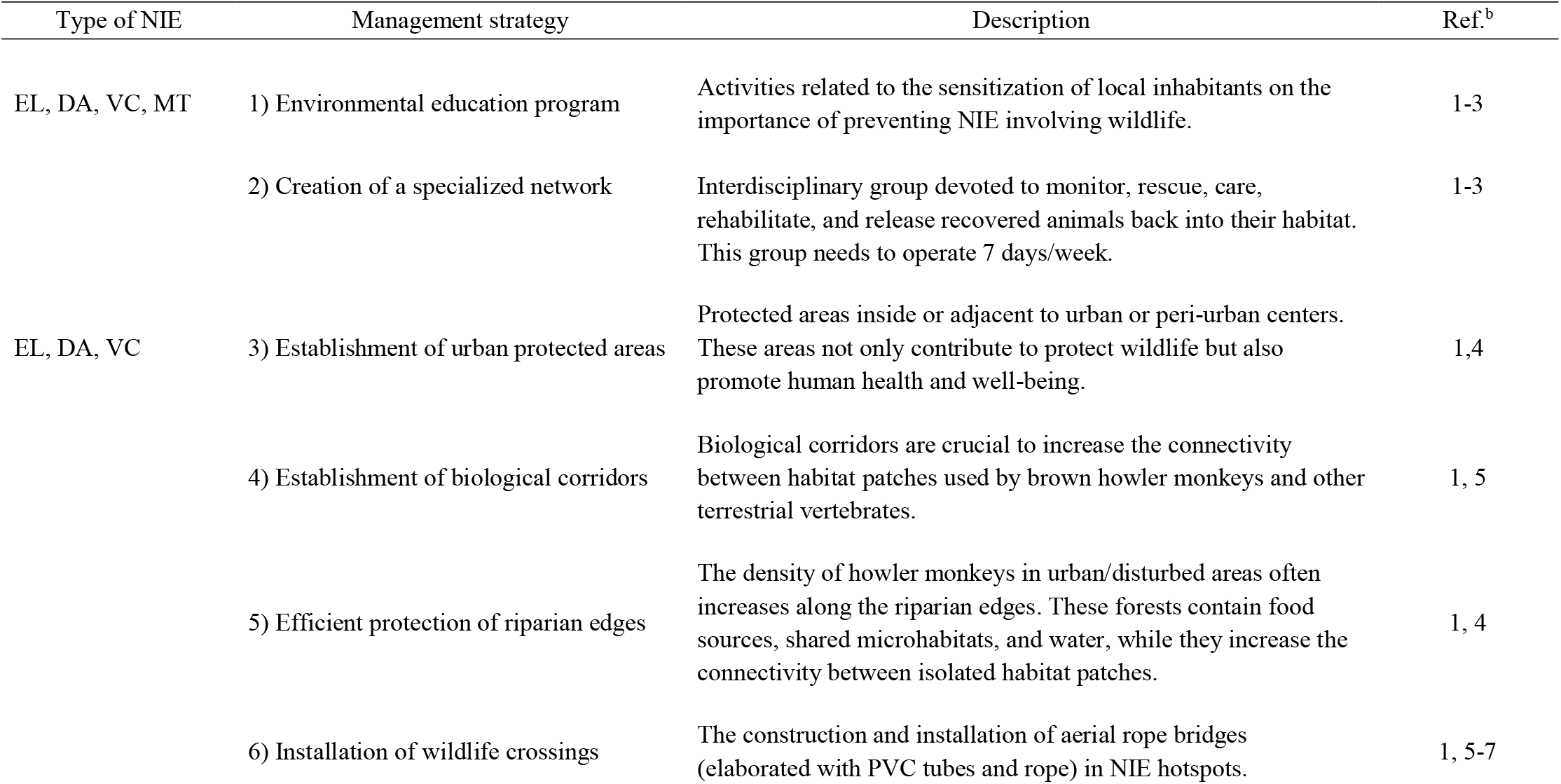
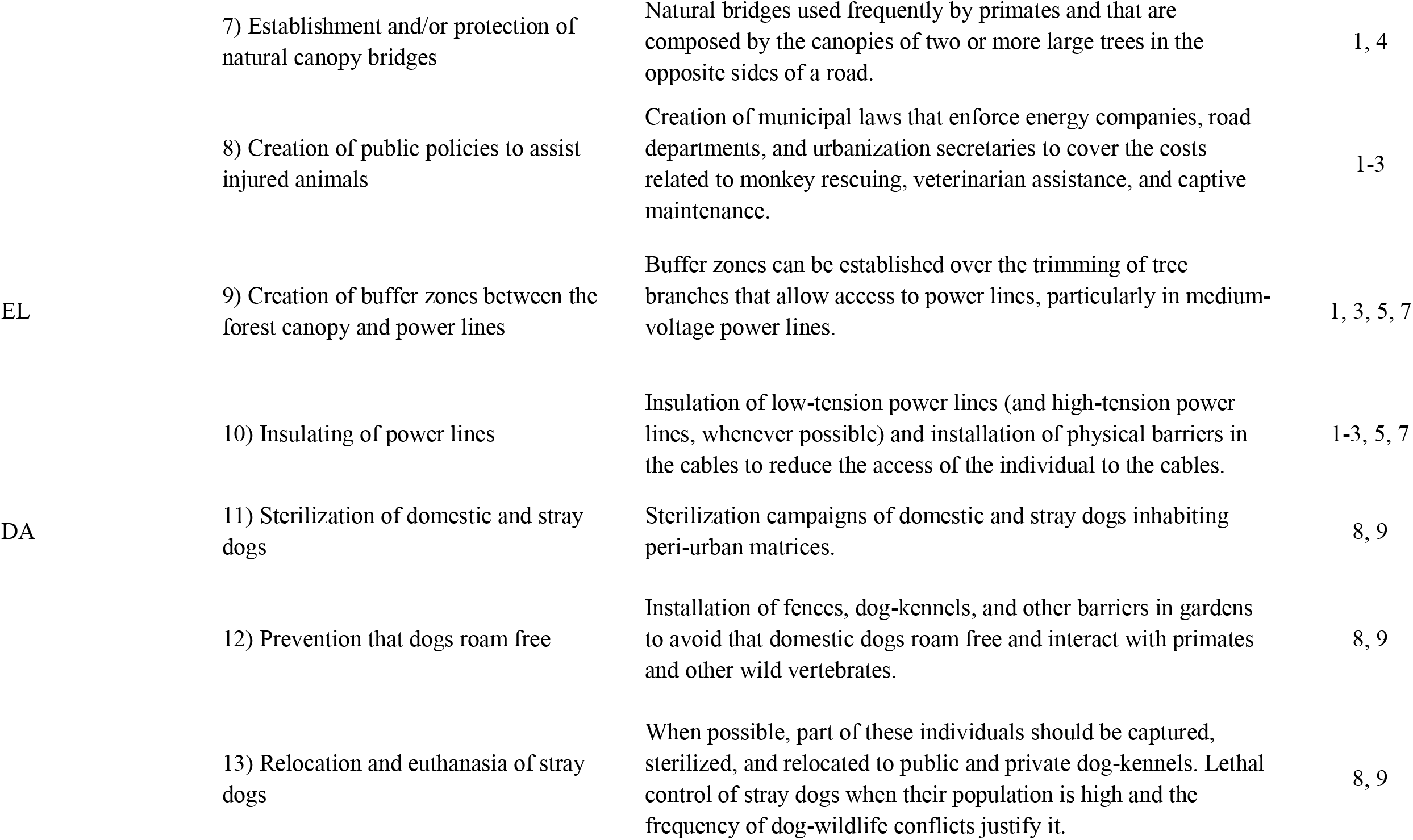
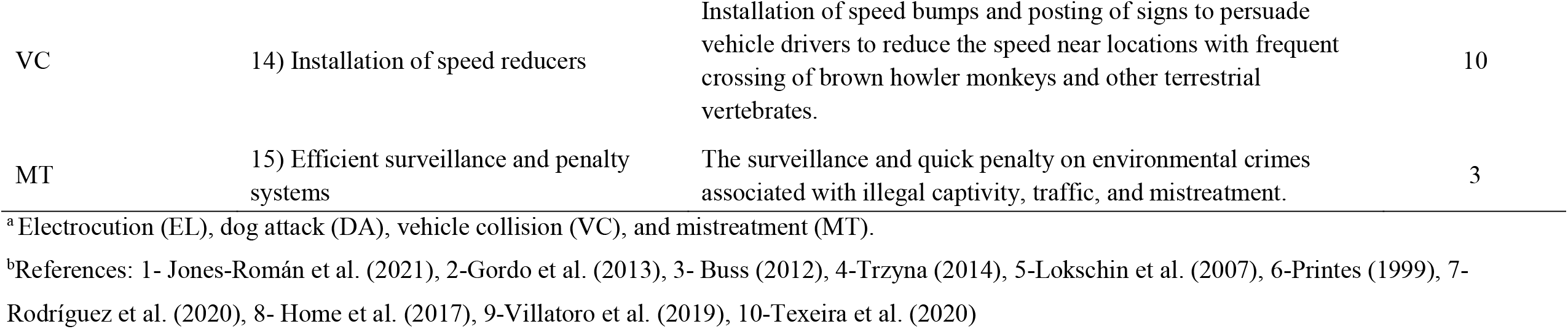
Fifteen potential management strategies to minimize the number of conflicts involving brown howler monkeys in anthropogenic peri-urban matrices of south Brazil

Attacks by stray or domestic dogs were the second major incident involving brown howler monkeys in both study regions. This finding supports evidence that domestic and feral dogs regularly kill primates (e.g. *Alouatta guariba clamitans*: Buss, 2012; Bicca-Marques et al., 2020; Lopes et al., 2021; *C. nigritus*: Oliveira et al., 2008) and other terrestrial mammals (e.g. Buttler et al., 2004, Lacerda et al., 2009; Home et al., 2017; Gatti et al., 2018). In the study regions, these attacks often occur when brown howler monkeys descend to the ground to access cultivated fruits in subsistence orchards guarded by domestic dogs (Buss 2012; Chaves and Bicca-Marques, 2017; Corrêa et al., 2018) or, when they cross roads, gardens or pastures to access another Atlantic Forest remnant (Óscar M. Chaves and João Claudio Godoy, personal observations). In most cases, death or severe injuries (e.g. organ perforations, mutilations, multiple bites, and skin cuts: see database in Chaves et al., 2021) are the outcome of dog attacks. Critically injured individuals cannot be returned to their habitats. For instance, most (ca. 70%) brown howler monkeys surviving dog attacks in RS and sent to the Rincão do Araticum Wildlife Rescue Center for recovery have never returned to their habitats because of, mainly lung, infections or tail, foot, or hand amputations (Silvia B. Ribeiro, personal communication). Therefore, population control of stray dogs is a necessary management strategy to reduce dog-wildlife NIEs in the study peri-urban matrices (Table 4).

Although less frequently reported, vehicle collisions and human mistreatments also deteriorate the health and compromise the survival of brown howler monkeys in the study regions. These NIEs were expected given (i) the high fragmentation and urban encroachment into the Atlantic Forest remnants that brown howler monkeys inhabit (Ribeiro et al., 2009), (ii) the frequent use of the ground by brown howler monkeys that supplement their diets with wild and cultivated foods found in scattered food patches separated by roads and other potentially lethal landscape elements (Buss, 2012; Chaves and Bicca-Marques, 2017; Corrêa et al., 2018), as well as by those dispersing from their natal groups (Strier et al., 2001) isolated in the fragmented landscape, (iii) the howlers’ limited ability to travel fast on the ground, and (iv) the inefficient Brazilian public policies to prevent/mitigate road kills (Gonçalves et al., 2018). Vehicle collision is a major cause of wildlife mortality in southern Brazil (Teixeira et al., 2020). A country-wide estimate indicated that ca. 1.3 million vertebrates (10% of which are large/medium birds, reptiles, primates, and terrestrial mammals) are killed every day along the Brazilian network of streets and roads (CBEE, 2019). Despite the lack of reliable data on the number of primates affected by VC in Brazil each year, the country is considered a world leader in the frequency of primate roadkills together with Indonesia and Equatorial Guinea (Galea and Humle, 2021). Finally, the reported percentage of mistreatments is probably underestimated. This NIE is rarely denounced by local inhabitants probably because they are either afraid of retaliations or because they are poorly informed on how to fill out a complaint.

The longer arms reach of adults compared with that of immature individuals increases their risk of touching cables with opposite charges simultaneously (Printes, 1999), thereby potentially explaining the highest frequency of electrocuted adult brown howler monkeys. The greater number of dog attacks and vehicle collisions on adults is compatible with their leading role in group travel both on the ground (Bicca-Marques and Calegaro-Marques, 1997) and in the canopy, as reported for black-and-gold howler monkeys (*Alouatta caraya*; Fernandéz et al., 2013). Whereas these morphological and behavioral age differences may explain the prevalence of adults involved in the most common NIEs, two non-mutually exclusive hypotheses may explain the higher impact of MT on infant and juvenile individuals. First, these individuals may be orphans rescued or harassed by local people following the aforementioned NIEs (see Chaves et al., 2020). Second, the stressful conditions that characterize their captive maintenance as MT are incompatible with their survival to adulthood.

Currently, we cannot evaluate whether the patterns that we found on the impact of each type of NIE on brown howler monkey populations have been biased by potential differences in detectability or in the propensity of local people to report them. This uncertainty results from the wealth of interacting and confounding variables in opportunistic studies relying on citizen science such as ours. Therefore, addressing such complex challenge in large study regions will require herculean efforts to systematically monitor target animal populations using appropriately-designed methods and the full collaboration of local people.

Irrespective of NIE type, most brown howler monkeys suffered lethal or mild-medium injuries (e.g. lung perforations, severe skin-burns, and body mutilations; see Fig. S2) and died soon after the incident or a few hours later. The overall fate of brown howler monkeys involved in NIEs was even worse if we consider that 93 to 98% of those surviving following veterinarian care were condemned to captive life. Also, it is not possible to impede that the rare individuals returning to their habitat continue exposed to the same risks, as seen in handicapped baboons (*Papio ursinus*) that exploit cultivated fruit and human-provided foods in Cape Peninsula, South Africa (Beamish and O’Riain, 2014). These findings highlight that brown howler monkeys are in great danger in urban and peri-urban areas of southern Brazil as have been suggested for the study areas (e.g. Printes, 1999; Buss, 2012; Corrêa et al., 2018; Bicca-Marques et al., 2020). Victims of mistreatments were the exception. Probably owing to their use as pets, severe injuries were rare. It is not uncommon for “owners” to seek some veterinary care for their pets in the study regions (Gerson Buss and Júlio César Souza Jr., personal communication). However, as hypothesized above, the rarity of adult brown howler monkeys as pets places doubt on their long-term survival under these conditions.

The removal of individuals from wild populations via death or life in captivity compromises the long-term conservation of brown howler monkeys in Atlantic Forest fragments with cascading consequences at the community level given their important role as seed dispersers (Martins, 2006; Chaves et al., 2018). A single adult brown howler monkey can disperse ca. 52,000 >2-mm seeds per year (Chaves et al., 2018). Considering estimates of brown howler monkey population density in other areas of the state of RS (see Table S3), we estimate that the injured individuals that we have reported (*n* = 248) represent ca. 10% of the taxon’s population in the study region. Given the limitations of our data collection and the fact that adult females involved in NIEs can be pregnant, the estimate above is conservative.

We also need to consider the economic cost associated with the rescue, veterinary care, and maintenance of handicapped individuals. Overall, the rehabilitation of urban wildlife concerns managers in developing countries, such as Brazil, because its high cost is rarely reimbursed by local or state governments (Karesh, 1995; Perry et al., 2020). For instance, the costs associated with rehabilitation and maintenance of brown howler monkeys in RS and SC (considering the maximum lifespan reported for captive howler monkeys, i.e. 20 years) can reach US$45,000 per individual during a 15-year period (Table S4). Considering only the basic costs of maintenance in captivity, rescue centers may spend US$ 177/individual/month in SC (Júlio César Souza Jr., personal communication).

Despite evidence of seasonal patterns in the occurrence of NIEs (i.e. mostly in the summer, when there is an increase in tourism in the RS study region; Buss, 2012), we did not find consistent patterns in both study regions. NIEs occurred throughout the year in both RS and SC. However, whereas they occurred at higher frequencies in the summer and fall than in the winter in RS, we found no seasonal differences in NIE frequency in SC. Whether this difference simply reflects the temporal characteristics of the sampling efforts in RS and SC (see Methods) or legitimate differences in landscape use resulting from higher numbers of people living or visiting the RS study region or driving through it during their summer vacation remains to be investigated. In SC, the sampling conducted by Projeto Bugio-FURB (https://www.furb.br/web/5579/projeto-bugio/apresentacao) was more uniformly distributed throughout the year. The leading role of Projeto Bugio, a research institute with a consolidated history of rescuing, caring, and rehabilitating brown howler monkeys in SC, may also explain the marked influence of study region on NIE-related mortality (Table 4).

## 5. Conclusions

To the best of our knowledge, this is the first study collating long-term data on the main threats faced by wild Neotropical primates living in peri-urban landscapes. We confirmed the aforementioned negative impacts of urbanization on wildlife health and survival described in short-term studies of primates and other vertebrates. We found that the fragmentation and urbanization of the Atlantic Forest represent serious (and often ignored) conservation challenges for the long-term survival of arboreal primates (and probably many other vertebrates). The severity of this scenario is further highlighted by the fact that despite flexibly adjusting their behavior to diverse anthropogenic landscapes (including peri-urban regions), alike other vertebrates (Sol et al., 2010; Schell et al., 2020), the long-term persistence of howler monkeys (*Alouatta* spp.) in fragmented peri-urban landscapes is at high risk (Bicca-Marques et al., 2020). As we have shown conservatively, hostile elements of the urban matrix, such as power lines, roads, domestic dogs, and wildlife traffickers, impose a much higher death rate to peri-urban populations than that seen in habitats more isolated from people. Therefore, designing and implementing appropriate strategies to prevent or mitigate human-wildlife NIEs are crucial to save urban-and peri-urban-tolerant species from extirpation.

## Acknowledgments

This study was supported by a grant from the Programa Nacional de Pós-Doutorado of the Coordenação de Aperfeiçoamento de Pessoal de Nível Superior – Brazil (Brazilian Higher Education Authority)/CAPES (Finance Code 001; PNPD grant # 2755/2010). JCBM thanks the Conselho Nacional de Desenvolvimento Científico e Tecnológico – Brazil (Brazilian National Research Council)/CNPq (PQ research fellowships #303154/2009-8, 303306/2013-0 and 304475/2018-1). We thank Dayse A. Rocha and Maria Carmen (Secretaria do Ambiente e do Desenvolvimento Sustentável/SEMA-RS), Silvia B. Ribeiro (Rincão do Aracutím), and many local inhabitants of the communities of Itapuã, Lami, Viamão, Blumenau, Indaial, and Pomerode, and to the Polícia Militar Ambiental of SC for the invaluable collaboration during the report of NIEs.

## Appendix A. Supplementary data

Supplementary data to this article can be found online at https://doi.org/

## Credit authorship contribution statement

**ÓMC:** Investigation, Conceptualization, Data collection, Data curation, Formal analysis, Writing original draft, Manuscript revision. **JCSJ:** Data collection, Data curation, Manuscript revision. **GB:** Data collection, Data curation, Manuscript revision, **ZMBH:** Data curation, Funding acquisition, Manuscript revision. **MMAJ:** Data collection, Data curation, Methodology, Manuscript revision. **ELSA:** Data curation, Methodology, Manuscript revision. **JCG:** Data collection, Methodology. **ARP:** Data collection.**TM:** Data collection, Data curation. **JCBM:** Funding acquisition, Supervision, Methodology, Manuscript revision, English revision.

