## Supplementary Material for "Wildlife is imperiled in peri-urban landscapes: threats to arboreal mammals"

**Table S1.** Municipalities and districts with records of negative interaction events (NIEs) involving brown howler monkeys in peri-urban matrices in Rio Grande do Sul and Santa Catarina states, southern Brazil.

| **State** | **Municipality** | **District/county** | **# NIEs** | **%TNC** |
| --- | --- | --- | --- | --- |
| Rio Grande do Sul | Viamão | Viamão | 90 | 36.3 |
|  | Porto Alegre | Lami | 83 | 33.5 |
|  | Viamão | Itapua | 28 | 11.3 |
|  | Porto Alegre | Porto Alegre | 9 | 3.6 |
|  | Porto Alegre | Canta Galo | 7 | 2.8 |
|  | Porto Alegre | Belem Novo | 5 | 2.0 |
|  | Porto Alegre | Restinga Nova | 6 | 2.4 |
|  | Porto Alegre | Lageado | 4 | 1.6 |
|  | Porto Alegre | Morro da Extrema | 3 | 1.2 |
|  | Porto Alegre | Santa Maria | 2 | 0.8 |
|  | Venâncio Aires | Venâncio Aires | 2 | 0.8 |
|  | Barra do Ribeiro | Barra do Ribeiro | 1 | 0.4 |
|  | Eldorado do Sul | Eldorado do Sul | 1 | 0.4 |
|  | Gravataí | Gravataí | 1 | 0.4 |
|  | Igrejinha | Igrejinha | 1 | 0.4 |
|  | Ivoti | Ivoti | 1 | 0.4 |
|  | Osório | Osório | 1 | 0.4 |
|  | São Leopoldo | São Leopoldo | 1 | 0.4 |
|  | Sapiranga | Sapiranga | 1 | 0.4 |
|  | Sarandi | Sarandi | 1 | 0.4 |
|  | **∑ = 12** | **20** | **248** | **100** |
| Santa Catarina | Blumenau | Blumenau | 150 | 51.5 |
|  | Indaial | Indaial | 60 | 20.6 |
|  | Pomerode | Pomerode | 25 | 8.6 |
|  | Jaraguá do Sul | Jaraguá do Sul | 7 | 2.4 |
|  | Timbó | Timbó | 5 | 1.7 |
|  | Florianópolis | Florianópolis | 4 | 1.4 |
|  | Gaspar | Gaspar | 4 | 1.4 |
|  | Rodeio | Rodeio | 4 | 1.4 |
|  | Ascurra | Ascurra | 3 | 1.0 |
|  | Lages | Lages | 3 | 1.0 |
|  | Laguna | Laguna | 3 | 1.0 |
|  | Brusque | Brusque | 2 | 0.7 |
|  | Campo Alegre | Campo Alegre | 2 | 0.7 |
|  | Massaranduba | Massaranduba | 2 | 0.7 |
|  | Rio dos Cedros | Rio dos Cedros | 2 | 0.7 |
|  | São Bento do Sul | São Bento do Sul | 2 | 0.7 |
|  | Antonio Carlos | Antonio Carlos | 1 | 0.3 |
|  | Braço do Norte | Braço do Norte | 1 | 0.3 |
|  | Garuva | Garuva | 1 | 0.3 |
|  | Guabiruba | Guabiruba | 1 | 0.3 |
|  | Ibirama | Ibirama | 1 | 0.3 |
|  | Ilhota | Ilhota | 1 | 0.3 |
|  | Itajaí | Itajaí | 1 | 0.3 |
|  | Joinville | Joinville | 1 | 0.3 |
|  | Papanduva | Papanduva | 1 | 0.3 |
|  | Rio do Sul | Rio do Sul | 1 | 0.3 |
|  | São Bento do Sul | São Bento do Sul | 1 | 0.3 |
|  | São Bonifácio | São Bonifácio | 1 | 0.3 |
|  | Taio | Taio | 1 | 0.3 |
|  | **∑ = 29** | **29** | **291** | **100** |

^a^ Total number of NIEs involving brown howler monkeys (i.e. electrocution, vehicle collison, dog attack, and mistreatment, see Methods). Data ranked in decreasing order of the percentage of the total number of NIEs (%TNC) and, then, alphabetically.

**Table S2.** Number of brown howler monkeys that died following mild-medium, severe and lethal injuries associated with the four main NIEs reported in the two study regions in southern Brazil.

| **NIE** | **Rio Grande do Sul** | **Santa Catarina** | **Total** |
| --- | --- | --- | --- |
| Electrocution | 56 | 49 | 105 |
| Dog attack | 48 | 60 | 108 |
| Vehicle collision | 28 | 18 | 46 |
| Human mistreatment | 4 | 6 | 10 |
| Total | 136 | 133 | 269 |
| **%Total NIEs*** | **61** | **51** | **56** |

*****Percentage of the total number of negative interaction events with complete data reported in each study region (RS=222, SC=262, *N*= 484)

**Table S3.** Studies assessing the population density of brown howler monkeys in Rio Grande do Sul and Santa Catarina states, southern Brazil.

| **Study site^a^** | **Year** | **Size (ha)^b^** | **Demography indicators^c^** | | **Methodology** | **Effort^d^** | **Ref.^e^** |
| --- | --- | --- | --- | --- | --- | --- | --- |
|  |  |  | D (ind./ha) | Pop. (ind.) |  |  |  |
| *Rio Grande do Sul* |  |  |  |  |  |  |  |
| Morro São Pedro | 2015 | 1,200 | 1.4 | 1,680 | Linear transects | 6 (205 km) | 1 |
| Reserva Econsciência | 2004 | 80 | 1.5 (0.7-1.9) | 120 | Linear transects | 1.5 (35 km) | 2 |
| Morro da Extrema | 1999 | 27 | 1.1 | 30 (415) | Grid cell counts | 1 (1.1 ha) | 3 |
| Morro da Extrema | 2001 | 86 | 1 | 86 | Grid cell counts | 13 (20.3 ha) | 4 |
| Mata de restinga, Recanto do lago, Lami | 1999 | 12.5 | 1.6 | 20 | Grid cell counts | 1 (0.4 ha) | 3 |
| Mata de restinga, Recanto do Lago, Lami | 2001 | 14 | 2,6 | 36 | Grid cell counts | 13 (7.3 ha) | 4 |
| Reserva Biológica do Lami | 2020 | 204 | — | >46 | Grid cell counts | 18 (54 ha) | 5 |
| Morro do Campista, PEI | 2000 | 284 | 0.8 (0.7-0.9) | 227 | Linear transects | 8 (107 km) | 6 |
| Morro Fortaleza, PEI | 2001 | 171 | 1,1 | 188 | Grid cell counts | 13 (16 ha) | 4 |
| Morro do Araça, PEI | 1994 | — | 0.9 | — | Grid cell counts | — | 7 |
| *Santa Catarina* |  |  |  |  |  |  |  |
| Morro Geisler | 2009 | 30 | 1.5 | 45 | Linear transects | 5 (167 km) | 8 |

^a^ Reserva Econsciência is part of Morro São Pedro, the largest Atlantic Forest fragment of Porto Alegre municipality. Morro da Extrema is located ca. 2 km from Morro São Pedro and 5 km from Mata de restinga, Lami. Morro Geisler (26º53`42”S, 49º13´34”W) is part of Parque Nacional da Serra do Itajaí, a secondary protected forest in the Itajaí municipality.

^b^ Size of the study site as reported by the authors. In sites >80 ha, this area included Atlantic Forest patches and other types of land cover (e.g. natural grasslands or ‘campos rupestres’, unpaved roads, small human settlements or isolated houses, and small agricultural fields).

^c^ D= density of individuals. Pop.= population density. In each study, population density was estimated by multiplying the mean density of individuals by the reported size of the study site in hectares. The number in parentheses for Morro da Extrema represents the estimate of the total population considering the total Atlantic Forest cover in the site (i.e. 377 ha, see Fialho 2000). — Data no available.

^d^ Sampling effort. The number of sampling months is indicated. In parentheses the number of kilometers walked or the total size of the area monitored.

^f^ References: **1**= Camaratta et al. (2017), **2**= Alonso (2004), **3**= Fialho (2000), **4**= Jardim (2005), **5**= Alfaya et. al (in press), **6**= Buss (2001), **7**= Cunha (1994), **8**= Junglos et al. (2009). With the exception of study 1, the other studies are unpublished (i.e. dissertations or monographs). Studies 1-3 used linear transect counts combined following the 'Distance sampling' method (Buckland et al., 2010), while study 6 was based only on linear transect counts.

**Table S4.** Estimate of financial costs associated with the rescue, veterinary care and maintenance of injured brown howler monkeys in wildlife rescue centers in Rio Grande do Sul (RS) and Santa Catarina (SC) states, southern Brazil.

| **Cost type** | **Description** | **Financial costs (US$)** | | **Source^a^** |
| --- | --- | --- | --- | --- |
|  |  | RS | SC |  |
| *I. Howler-related cost* |  |  |  |  |
| Rescue of injured animal(s) | Minimal costs related to rescuing injured monkeys on roads, powerlines or rural gardens | 200 | ̶ | 1,2 |
| Veterinarian emergency treatment | Veterinarian emergency care of monkeys with medium to severe injuries (including surgeries) | 300 | ̶ | 3 |
| Basic veterinarian care/year | Routine veterinarian care of monkeys in captivity, including antibiotics, nutritional supplements, and antiparasitic drugs | 100 | 100 | 3 |
| Feeding/year | Cost associated with feeding injured/crippled monkeys unable to return to the wild | 2130 | 2142 | 4 |
| *II. Legal costs* |  |  |  |  |
| Potential legal penalties/year | Penalty fees established in the Brazilian Environmental Laws (Lei de Crimes Ambientais No. 9.605/98) for companies responsible for road construction or energy supply due to law violations | ̶ | 475,312* | 5 |
| **Total costs/first year/monkey** |  | 2730 | 2242 |  |
| **Total costs/subsequent years** |  | 2230 | 2242 |  |
| **Total costs/monkey/15 years** |  | **45,100** | **44,840** |  |

^a^Source of information: **1**= OMC (personal observation), **2**=JCG (personal observation), **3**= Silvia B. Ribeiro, Mantenedouro de Fauna Rincão do Araticum (personal communication), **4**= Parecer No. MA-1665-2011, Ministério Público do Rio Grande do Sul (<https://doi.org/10.17632/wv8pvzwfyw.1>), **5**= Proyecto Fauna Viva, Florianópolis, SC (personal communication)

*This estimate was based on the mean number of wild monkeys electrocuted in the municipality of Blumenau, Santa Catarina, per year.

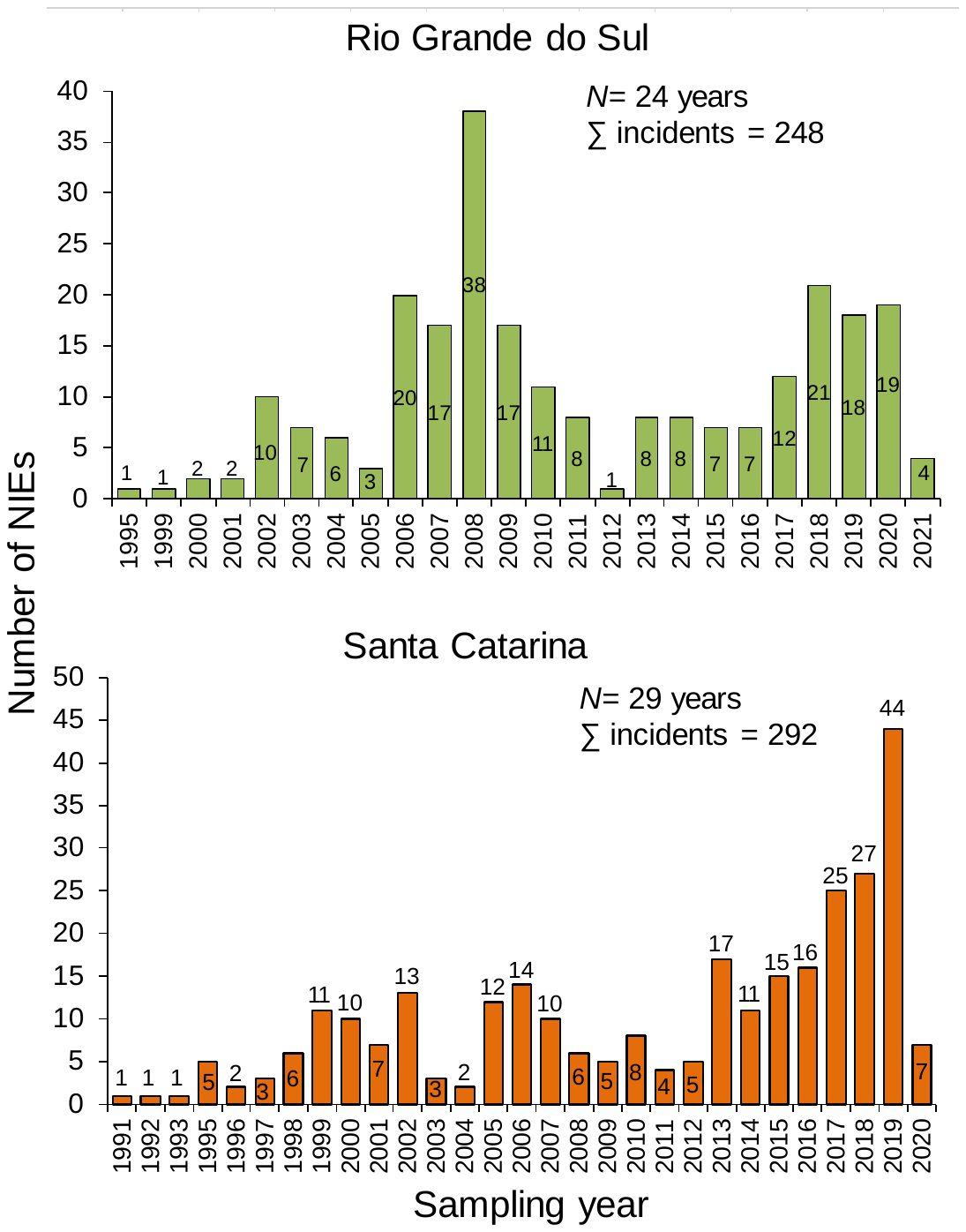

**Fig. S1.** Annual number of NIEs involving brown howler monkeys in urban and peri-urban areas of Rio Grande do Sul and Santa Catarina states (*N=* 540 NIEs). Further details in Methods.

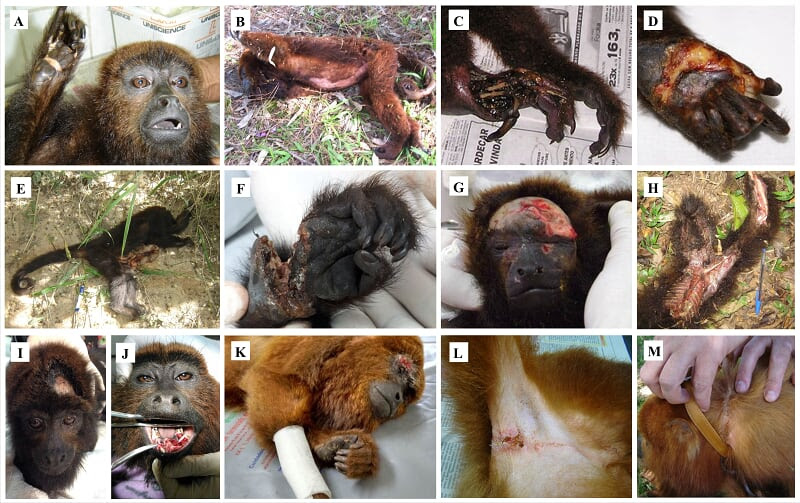

**Fig. S2.** Main types of external injuries found in brown howler monkeys (*Alouatta guariba clamitans*) victims of electrocution, vehicle collision, dog attack and mistreatment in urban and peri-urban areas in southern Brazil. Skin-burns and mutilations in hands and toes resulting from electrocution in power lines (A-D); lethal injuries resulting from collision with a vehicle in the road Frei Pacifico ERS-118 (E, F); severe eye and skin injuries after dog attacks (G, H); victims of mistreatment by local inhabitants: air-gun pellet injured in the head (I), mouth hematoma (J), broken arm and eye hematoma (K); chest and neck injuries in an adult male victim of illegal captivity (L, M). Photos by Thais Michel (A, L), João Claudio Godoy (B, D), Urban Monkey Program (UFRGS) (C), Gerson Buss (E, H, M), Renan Stadler (F, K), Elisandro Santos (G), Priscila Medina, Preservas (UFRGS) (I), and Bruna Zafalon (J).
